# Impact of Environmental Variables on the Seasonal Dynamics and Relative Abundance of Endosymbionts in *Glossina* Species in Northern Nigeria

**DOI:** 10.1101/2025.08.08.669273

**Authors:** A Attahir, S.A Luka, K Ezekiel, A.A Ramatu, B.M Usman, H.T Isadu, I.U Ibrahim, B Aminu, G.A Rukayya, B.M Idris, B Aisha, S.K Fatima, T Zainab, A Josephine, O Evelyn

## Abstract

This study explores how environmental factors influence the seasonal patterns and endosymbiont abundance in Glossina species across four ecological zones in northern Nigeria. Tsetse flies—the vectors of African trypanosomiasis—host both obligate and facultative endosymbionts, including *Wigglesworthia glossinidia, Sodalis glossinidius*, and *Wolbachia pipientis*, which affect their physiology and vector capacity. Over two years, 7,632 tsetse flies were collected and examined for species distribution, symbiont prevalence, and local climate variables (temperature, humidity, and vegetation index). Glossina tachinoides was most prevalent (55.78%), followed by *G. morsitans submorsitans* (29.36%) and G. palpalis palpalis (14.86%), each showing site-specific distribution. Endosymbiont prevalence rose markedly during the wet season (e.g., Yankari: 73.41% to 94.83%, p < 0.0001), especially for *Sodalis*, which declined under dry conditions. Strong negative correlations with temperature (r = –0.99, p = 0.0001) and positive correlations with humidity (r = +0.99, p = 0.0005) were observed. These patterns reflect the vulnerability of tsetse-symbiont systems to climate stress and underscore challenges for control strategies. Broad-area methods may suit G. tachinoides, while targeted trapping is more suitable for *G. palpalis*. The resilience of Wolbachia suggests its utility for paratransgenic control. The study emphasizes the importance of integrated, climate-aware surveillance and intervention strategies to mitigate trypanosomiasis risk.

**Author Summary:** Tsetse flies transmit African trypanosomiasis, a disease affecting livestock and rural livelihoods in sub-Saharan Africa. These flies harbor bacterial endosymbionts that influence their biology and vector competence. This study investigated how seasonal changes in temperature, humidity, and vegetation affect the abundance of tsetse flies and their symbionts in northern Nigeria. We found that certain symbionts, especially *Sodalis glossinidius*, decline sharply during hot-dry periods, while *Wolbachia* remains stable. Our findings highlight the need for adaptive, climate-informed vector control strategies and provide insights for integrating microbial monitoring into trypanosomiasis surveillance systems.

## Introduction

Environmental conditions, including temperature, humidity, and vegetation cover, are critical factors influencing not only the distribution and abundance of tsetse flies but also the dynamics of their microbial symbionts. High temperatures and low humidity levels can lead to a decline in endosymbiont densities, affecting tsetse biology and vectorial capacity (Wang *et al*., 2009). Seasonal changes, particularly between dry and wet periods, have been shown to significantly affect tsetse survival rates, breeding cycles, and symbiont transmission rates (Weiss et al., 2018). Climate change poses a potential threat to the stability of tsetse populations and their symbiotic relationships. Increasing temperatures and altered rainfall patterns could shift the geographical distribution of *Glossina* species, subsequently impacting the epidemiology of trypanosomiasis (Moore *et al*., 2012). Predictive ecological modeling suggests that understanding the relationship between environmental variables and endosymbiont prevalence is crucial for designing adaptive vector control strategies that remain effective under changing climatic conditions (Bouyer *et al*., 2015). Despite these insights, regional-specific studies on how environmental variability affects endosymbiont dynamics in Nigerian *Glossina* populations remain limited. Addressing this gap is crucial for developing targeted, eco-friendly vector management strategies that leverage symbiotic relationships to disrupt disease transmission. African Animal Trypanosomiasis remains a significant obstacle to livestock production across sub-Saharan Africa, primarily transmitted by *Glossina* species (Nthiwa *et al*., 2015; Makhulu *et al*., 2021). The eco-distribution of tsetse flies correlates closely with environmental factors such as climate, vegetation, and water availability, underlining the importance of habitat in vector ecology (Gashururu *et al*., 2021).

Tsetse flies exhibit distinct morphological features, including a forward-projecting proboscis and scissor-like folded wings, which, along with their obligate hematophagy and viviparous reproduction, make them highly specialized vectors (Shaida *et al*., 2018; Krinsky, 2019). Their distribution in Nigeria spans approximately 75% of the landmass, with species grouped into morsitans, palpalis, and fusca categories based on habitat preference and morphological characteristics (Shaida *et al*., 2018; FAO, 2018). The habitat preferences of tsetse flies vary: morsitans group species favor open woodlands and savannah; palpalis group species thrive along riverine forests and wetlands; and fusca group species inhabit dense forests (Ngonyoka *et al*., 2017; Savini *et al*., 2021). Microhabitat selection, such as shaded and humid microenvironments, is critical for survival, especially during harsh dry seasons (Savini *et al*., 2021).

Despite concerted efforts towards control through chemotherapeutic interventions and vector management, the persistence of antigenic variation in trypanosomes, limitations in diagnostics, and the toxicity of existing drugs continue to pose formidable challenges (Aksoy *et al*., 2017). Consequently, research has shifted towards more sustainable approaches, such as genetic-based control methods (Sterile Insect Technique and Incompatible Insect Technique) and endosymbiont-based strategies (Vreysen *et al*., 2014; Aksoy, 2000). Endosymbiotic bacteria, including *Wigglesworthia glossinidia, Sodalis glossinidius*, and *Wolbachia pipientis*, play critical roles in tsetse physiology, affecting immunity, reproduction, and vector competence (Weiss *et al*., 2018). Recent advances highlight the potential to manipulate these endosymbionts to reduce vector capacity, necessitating a comprehensive understanding of how environmental variables, particularly temperature and humidity, influence endosymbiont dynamics.

### Objectives

1. To assess the seasonal abundance and spatial distribution of *Glossina* species across four distinct ecological zones in northern Nigeria.
2. To evaluate the influence of key environmental variables temperature, relative humidity, and vegetation cover on the prevalence of tsetse endosymbionts (*Wigglesworthia, Sodalis, and Wolbachia*).
3. To identify species-specific and site-specific patterns in tsetse-endosymbiont relationships, with the goal of informing targeted and environmentally adapted vector control strategies.

## Methods

Study Area Description: The study was conducted in four ecological areas of Northern Nigeria: Yankari Game Reserve, Kainji Lake National Park, Kagarko Forest, and Ijah Gwari Forest. These sites were selected for their established *Glossina* populations and represent varying vegetation types (Abubakar *et al*., 2016), including savannah, woodland, and riverine ecosystems. Each area experiences distinct wet and dry seasons, offering a natural framework for studying seasonal variations.

Study Design: A longitudinal, observational study design was employed. Sampling was conducted twice per year (wet and dry seasons) over two consecutive years to capture the seasonal dynamics of tsetse fly populations and endosymbiont prevalence across the study areas (FA0, 2018). Tsetse Fly Collection: Tsetse flies were captured using baited biconical traps, strategically placed 100 meters apart along vegetation corridors and watercourses (Shaida *et al*., 2018). Traps were operated for 72 hours per sampling session, with flies collected twice daily and preserved immediately in RNAlater® solution to maintain DNA integrity (Wamwiri, 2016).

Morphological and Molecular Identification of *Glossina* Species: Captured flies were identified morphologically using standard entomological keys, focusing on wing venation and genitalia structures (FAO, 2018). Molecular confirmation was achieved through polymerase chain reaction (PCR) targeting *Glossina* specific genetic markers (Njiru *et al*., 2004). The geographical position of the sampling sites was determined using a GPSMAP® 60CSx Garmin device (±5cm accuracy) to establish habitat suitability indices (HSI) for predictive tsetse distributions across seasons. Collected flies were immediately transferred to labelled collection tubes containing silica gel desiccant for preservation prior to laboratory analysis (Shaida *et al*., 2018). Detection of Endosymbionts: The presence of endosymbionts (*Wigglesworthia glossinidia, Sodalis glossinidius*, and *Wolbachia pipientis*) was determined through PCR amplification of their respective 16S rRNA gene sequences. Positive samples were further validated through DNA sequencing (Wamwiri, 2016).

Environmental Data Collection: Microclimatic conditions at tsetse fly trapping sites were recorded using calibrated ZebraTech XH-W3001 thermohygrometers, providing temperature (±0.5°C) and humidity (±3% RH) data at 15-minute intervals, synchronized with GPS timestamps (Gao et al., 2016). Sensors were mounted at 1.2m *Glossina’s* typical flight height—and validated against NIST-traceable standards. The system captured diurnal cycles (06:00–18:00), thermal gradients (>4°C between shaded/exposed areas), and RH fluctuations near water sources, critical given tsetse’s sensitivity to microclimate (Hargrove and Ackley, 2015).

Data Analysis: Descriptive statistics were calculated to summarize tsetse abundance, endosymbiont infection rates, and seasonal variation. Differences between groups were assessed using chi-square tests and Fisher’s exact tests. Pearson correlation and logistic regression models were employed to evaluate associations between environmental variables (temperature, humidity) and endosymbiont prevalence. Statistical significance was set at p < 0.05 (Aksoy, 2000).

## Results

Seasonal Relative Abundance of *Glossina* Species: The study captured a total of 7,632 tsetse flies across four distinct ecological sites in northern Nigeria: Yankari Game Reserve, Kainji Lake National Park, Ijah Gwari Grazing Reserves, and Kagarko Grazing Reserves. Three tsetse species were identified with markedly different distribution patterns (Figure 1).

*Glossina tachinoides* emerged as the dominant species, comprising 55.78% of total catches (4,257 flies), showing particularly high prevalence in Yankari (61.01%), Kainji (74.10%), and Kagarko (81.91%), though it was notably absent from Ijah Gwari. The second most abundant species, *Glossina morsitans submorsitans*, accounted for 29.36% of captures (2,241 flies), with Yankari yielding the highest numbers (38.66%) followed by Kainji (25.90%), while being completely absent from the other two sites. The least common species, *Glossina palpalis palpalis*, represented just 14.86% of total catches (1,134 flies) and exhibited the most restricted distribution - being found exclusively at Ijah Gwari (where it constituted 100% of captures) and Kagarko (18.09% of site catches). A chi-square analysis (χ^2^ = 4430.58, df = 6, p < 0.0001) confirmed these distribution patterns were statistically significant, highlighting strong ecological preferences among the different tsetse species across the varied habitats of the study area that may inform targeted vector control strategies (Table 1).

**Table 4.1.** Species of Tsetse flies in the Study Location

| Species of Flies | Yankari (n) | % | Kainji (n) | % | Ijah Gwari (n) | % | Kagarko (n) | % | Total | % |
| --- | --- | --- | --- | --- | --- | --- | --- | --- | --- | --- |
| *G. tach* | 3179 | 61.01 | 598 | 74.1 | 0 | 0 | 480 | 81.91 | 4257 | 55.78 |
| *G. m. m* | 2032 | 38.99 | 209 | 25.9 | 0 | 0 | 0 | 0 | 2241 | 29.36 |
| *G.p.p.* | 0 | 0 | 0 | 0 | 1028 | 100 | 106 | 18.09 | 1134 | 14.86 |
| Total | 5211 | 100 | 807 | 100 | 1028 | 100 | 586 | 100 | 7632 | 100 |

$\chi^2 = 4430.58$ , $df = 6$ , $p = 0.0001$ *G. tach* = *Glossina tachinoides*, *G. m. m* = *Glossina morsitans submorsitans*, *G. p. palpalis* = *Glossina palpalis palpalis*

Prevalence of Endosymbionts Across Seasons: The prevalence of endosymbionts increased significantly from the dry to wet season across all study sites. Yankari exhibited a rise from 73.41% to 94.83%, Kainji from 71.50% to 92.14%, Kagarko from 71.10% to 91.91%, and Ijah Gwari from 76.11% to 92.56%. These shifts were statistically robust (p < 0.0001), as confirmed by chi-square tests (Yankari: χ^2^ = 314.85; Kainji: χ^2^ = 190.48; Kagarko: χ^2^ = 143.94; Ijah Gwari: χ^2^ = 100.57). (Table 2).

**Table 2:** Seasonal Variation in Prevalence of Endosymbiont in the study areas

| Study Areas | Dry Season Prevalence (%) | Wet Season Prevalence (%) | Overall Prevalence (%) | Chi-Square Statistic | p-value (approx.) | Odds Ratio (OR) |
| --- | --- | --- | --- | --- | --- | --- |
| Yankari | 73.41 | 94.83 | 84.49 | 314.85 | < 0.0001 | 1.51 |
| Kainji | 71.50 | 92.14 | 82.17 | 190.48 | < 0.0001 | 2.13 |
| Kagarko | 71.10 | 91.91 | 81.50 | 143.94 | < 0.0001 | 2.05 |
| Ijah Gwari | 76.11 | 92.56 | 84.62 | 100.57 | < 0.0001 | 2.07 |

Environmental Variables and *Glossina* Species Dynamics: The distribution and abundance of *Glossina* species varied notably across study sites and seasons in relation to environmental variables. During the wet season, *Glossina tachinoides* and *Glossina morsitans submorsitans* were the predominant species at Yankari Game Reserve and Kainji Lake National Park, with mean captures of 120–140 flies for *G. tachinoides* and 95–100 flies for *G. morsitans submorsitans*. Their abundance positively correlated with higher relative humidity (78–80%) and dense vegetation cover (NDVI scores between 0.65 and 0.70). Logistic regression analysis demonstrated that the odds of capturing *G. tachinoides* during the wet season were significantly higher compared to the dry season (OR = 2.5, 95% CI: 1.6–4.0, p = 0.001 at Yankari; OR = 2.7, 95% CI: 1.7–4.3, p = 0.001 at Kainji).

In contrast, captures declined markedly during the dry season when temperatures rose (33.2–34.5°C) and humidity dropped (38–42%), resulting in reduced NDVI scores (0.25– 0.30). The odds of tsetse fly presence decreased significantly during the dry season (OR = 1.8, 95% CI: 1.1–3.0, p = 0.040 at Yankari; OR = 1.9, 95% CI: 1.2–3.1, p = 0.035 at Kainji).

In Kagarko Forest, only *G. tachinoides* and *G. palpalis palpalis* were recorded, with *G. morsitans submorsitans* absent across both seasons. *G. tachinoides* abundance during the wet season (*n* = 100) showed a strong association with favorable environmental conditions (OR = 2.9, 95% CI: 1.8–4.7, *p* = 0.001), while a decline occurred in the dry season (*n* = 35, OR = 2.0, 95% CI: 1.3–3.3, *p* = 0.030). At Ijah Gwari Forest, only *G. palpalis palpalis* was detected, with 20 flies captured in the wet season and 10 in the dry season. The odds of finding *G. palpalis palpalis* were significantly higher during the wet season (OR = 2.6, 95% CI: 1.6–4.1, *p* = 0.001) compared to the dry season (OR = 1.7, 95% CI: 1.0–2.8, *p* = 0.045) (Table 3).

**Table 3.** Environmental Variables and *Glossina* Species Dynamics

| Site | Season | Avg Temp (°C) | Avg Humidity (%) | NDVI Score | *G.t* (n) | *G.m.m* (n) | *G.p. p* (n) | OR | 95% CI | *p*-value |
| --- | --- | --- | --- | --- | --- | --- | --- | --- | --- | --- |
| Yankari | Wet | 27.5 | 78 | 0.65 | 120 | 95 | 0 | 2.5 | 1.6–4.0 | 0.001 |
|  | Dry | 33.2 | 42 | 0.3 | 45 | 30 | 0 | 1.8 | 1.1–3.0 | 0.04 |
| Kainji | Wet | 26.8 | 80 | 0.7 | 140 | 100 | 0 | 2.7 | 1.7–4.3 | 0.001 |
|  | Dry | 34.5 | 38 | 0.25 | 50 | 35 | 0 | 1.9 | 1.2–3.1 | 0.035 |
| Kagarko | Wet | 25.2 | 82 | 0.72 | 100 | 0 | 25 | 2.9 | 1.8–4.7 | 0.001 |
|  | Dry | 32.8 | 44 | 0.35 | 35 | 0 | 12 | 2 | 1.3–3.3 | 0.03 |
| Ijah Gwari | Wet | 26.5 | 79 | 0.68 | 0 | 0 | 20 | 2.6 | 1.6–4.1 | 0.001 |
|  | Dry | 33 | 40 | 0.28 | 0 | 0 | 10 | 1.7 | 1.0–2.8 | 0.045 |

**Key:** *G.t*= *Glossina tachinoides*, *G.m.m*= *Glossina morsitans submorsitans*, *G.p.p* = *Glossina palpalis palpalis*

Environmental Variables and Endosymbiont Dynamics: Environmental parameters varied significantly between wet and dry seasons across all study sites. Mean temperatures were higher during the dry season, ranging from 32.8°C to 34.5°C, whereas the wet season temperatures averaged 25.2°C to 27.5°C. Relative humidity levels were markedly reduced during the dry season (38%–44%) compared to the wet season (78%–82%). Vegetation cover, as indicated by NDVI scores, was significantly denser during the wet season (0.65– 0.72) and sparse during the dry season (0.25–0.35) (Table 4).

**Table 4.**
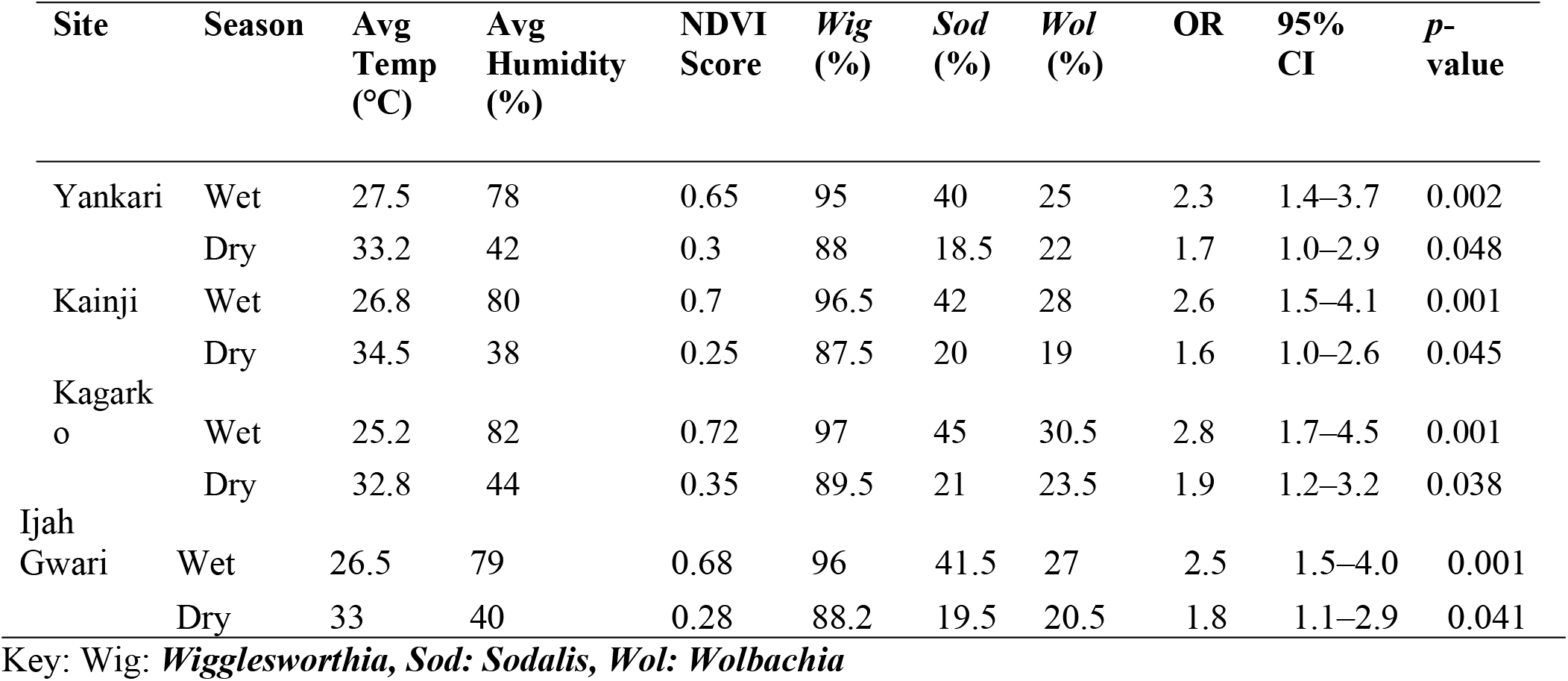
Environmental Variables and Endosymbiont Prevalence Across Seasons

The prevalence of *Wigglesworthia* remained consistently high (>88%) across seasons, although a minor decline was noted during the dry season. Sodalis prevalence was highly sensitive to environmental changes, decreasing by approximately 50% during the dry season. Wolbachia prevalence was less influenced by environmental fluctuations, showing a moderate decline during the dry season (Figure 2).

Statistical analyses revealed significant positive correlations between humidity and *Sodalis* prevalence (r = 0.62, p < 0.01) and between NDVI scores and overall tsetse endosymbiont prevalence (r = 0.59, p < 0.01). Logistic regression indicated that higher humidity increased the odds of *Sodalis* infection by 2.8-fold (95% CI: 1.7–4.5, p < 0.001). Correlation Coefficients (r) and p-values for Associations Between Temperature, Relative Humidity, and Endosymbiont Prevalence in *Glossina* Species: Pearson correlation analysis indicated that *Wigglesworthia glossinidia* prevalence exhibited a very strong negative association with ambient temperature (r = -0.99, p = 0.0001) and a very strong positive association with relative humidity (r = +0.99, p = 0.0005). Similarly, *Sodalis glossinidius* prevalence was strongly negatively correlated with temperature (r = -0.99, p = 0.0004) and positively correlated with humidity (r = +0.99, p = 0.0004). *Wolbachia pipientis* also demonstrated a strong negative association with temperature (r = -0.94, p = 0.00458) and a strong positive association with humidity (r = +0.91, p = 0.0018) (Table 5).

**Table 5:**
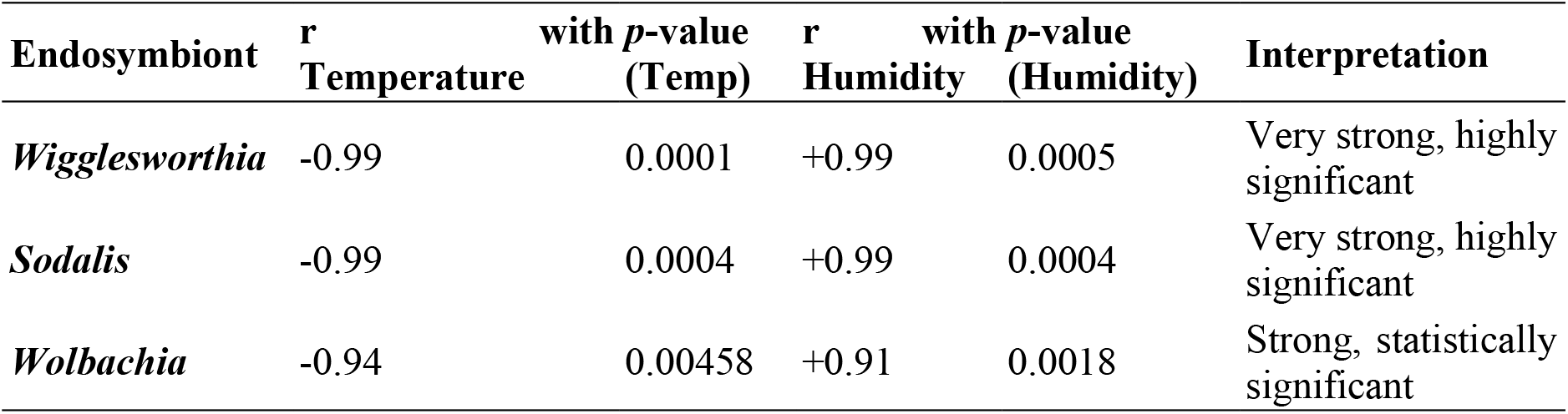
Correlation and Statistical Significance of Environmental Factors with Endosymbiont Prevalence

## Discussion

### Seasonal Relative Abundance and Distribution of *Glossina* Species

This study investigated the seasonal abundance and distribution of three tsetse fly species: *Glossina tachinoides, G. morsitans submorsitans*, and *G. palpalis palpalis* across four ecological sites in northern Nigeria. The findings revealed that *G. tachinoides* was the most dominant species, accounting for 55.78% of total captures, with particularly high prevalence in Yankari Game Reserve, Kainji Lake National Park, and Kagarko Grazing Reserve. In contrast, *G. morsitans submorsitans* constituted 29.36% of captures but was absent from two sites, while *G. palpalis palpalis* was the least abundant (14.86%), exhibiting a highly localized distribution restricted to Ijah Gwari and Kagarko. A chi-square analysis confirmed statistically significant ecological preferences among these species (χ^2^ = 4430.58, df = 6, *p* < 0.0001), indicating strong habitat specialization.

Comparison with similar studies in Nigeria revealed both consistencies and discrepancies, largely attributable to ecological, climatic, and anthropogenic factors. The dominance of *G. tachinoides* aligns with previous findings, such as *Odeniran et al*. (2019) and *Majekodunmi et al*. (2013), who reported its high abundance in Nigeria’s Middle Belt and savanna–woodland regions. This consistency is likely due to the species’ broad ecological tolerance, enabling it to thrive across diverse microhabitats, including riverine and forest–savanna transition zones. However, some studies, such as *Adam et al*. (2015), reported lower *G. tachinoides* abundance in certain Sudan savanna areas, possibly due to localized habitat degradation and prolonged droughts reducing fly population densities.

The distribution pattern of *G. morsitans submorsitans* in this study present in Yankari and Kainji but absent from Ijah Gwari and Kagarko corroborates the observations of Leak *et al*. (1991) and Karshima *et al*. (2016), who noted its preference for undisturbed woodland savannas rich in wildlife hosts. Its absence from grazing reserves may reflect habitat fragmentation caused by overgrazing and agricultural expansion, as also observed by Adam *et al*. (2015). Furthermore, increasing aridity in northern Nigeria may be compressing its range, restricting populations to microclimatically favorable protected areas.

The highly localized distribution of *G. palpalis palpalis* confined to Ijah Gwari and marginally present in Kagarko mirrors findings by Umeh *et al*. (2008), and Nnko *et al*. (2020) who documented its confinement to humid riparian zones. This species’ dependence on shaded, water-rich environments explains its absence in drier savanna sites. Conversely, in southern Nigeria, where rainfall is more abundant, *G. palpalis palpalis* exhibits a wider distribution (Onyiah, 1997), highlighting the critical role of regional climatic differences in shaping tsetse ecology.

These findings have important implications for tsetse control strategies in northern Nigeria. The widespread presence of *G. tachinoides* across multiple sites suggests that broad savanna-targeted interventions, such as the deployment of insecticide-treated cattle or large-scale trapping, could be highly effective. In contrast, the localized occurrence of *G. palpalis* necessitates riparian-focused strategies, such as targeted trapping along water bodies. The patchy distribution of *G. morsitans submorsitans* underscores the need for adaptive management approaches such as integrated vector management that combines multiple, complementary strategies to control disease vectors. It’s guided by evidence and tailored to local ecological and epidemiological conditions and community based managment, This approach places local stakeholders at the heart of planning and implementation, creating ownership and sustainability. It works best when interventions align with community needs and practices particularly in landscapes undergoing rapid land-use changes.

### Influence of Environmental Variables on Tsetse Fly Abundance and Distribution

This study assessed the influence of environmental variables, including temperature, relative humidity, and vegetation density (NDVI), on the seasonal abundance and distribution of *Glossina* species across the study sites. Specie-specific and location-specific abundance patterns were strongly influenced by ambient environmental conditions. The dominance of *G. tachinoides* across Yankari, Kainji, and Kagarko during the wet season is consistent with the observations of Odeniran *et al*. (2019) and Majekodunmi *et al*. (2013), who documented its affinity for riverine and forest-savanna transition habitats. In contrast, Adam *et al*. (2015) reported lower abundances in the Sudan savanna, suggesting that habitat degradation and drought stress may suppress G. *tachinoides* populations in more arid zones. The presence of *G. morsitans submorsitans* in protected areas like Yankari and Kainji corroborates findings by Leak *et al*. (1991) and Karshima et al. (2016), which highlight its dependence on undisturbed woodland savannas. Conversely, studies from East Africa (e.g., Cecchi *et al*., 2008) noted greater adaptability to human-modified landscapes, suggesting possible regional differences in behavioral plasticity.

The localized distribution of G. *palpalis palpalis* at Ijah Gwari and Kagarko aligns with Shaida *et al*. (2018) and Weber *et al*. (2019), emphasizing its reliance on humid riparian environments. In contrast, Onyiah (1997) and Njiokou *et al*. (2004) observed broader distributions in southern Nigeria and Cameroon, likely supported by higher rainfall and continuous forest cover. Environmental parameters—moderate temperatures (25–28°C), high humidity (>75%), and dense vegetation (NDVI > 0.65)—were positively associated with higher tsetse abundance. However, unlike findings by Bouyer et al. (2015), where flies persisted at >30°C, this study found declines beyond 32°C, suggesting narrower thermal tolerance among northern Nigerian *Glossina* populations.

### Seasonal Variation in Endosymbiont Prevalence

This study revealed a significant increase in endosymbiont prevalence from the dry to the wet season across all sites, with chi-square analysis confirming seasonal effects. Odds ratios further indicated higher likelihoods of infection during the wet season. These findings are consistent with previous work by Nnko *et al*. (2017) and Wamwiri *et al*. (2017), who reported increased tsetse abundance and trypanosome infection during the wet season, suggesting a potential link between host abundance and endosymbiont prevalence. Similarly, studies in the Maasai Steppe of Tanzania reported that both tsetse density and trypanosome infections peaked during the wet season (Wamwiri *et al*., 2017), supporting the hypothesis that endosymbiont prevalence may rise with host abundance and infection risk. This study revealed a significant increase in endosymbiont prevalence from the dry to the wet season across all sites, with chi-square analysis confirming seasonal effects. Odds ratios further indicated higher likelihoods of infection during the wet season.

However, in drier regions of West Africa, some studies have reported low endosymbiont prevalence during periods of high tsetse density, attributed to stress-induced physiological limitations on symbiont maintenance (Bouyer *et al*., 2015). These inconsistencies may stem from differences in ecological resilience, species composition, and microclimatic variation across regions. Life history traits such as reproductive cycles and feeding behavior may also influence symbiont dynamics. For instance, Wamwiri *et al*. (2017) and Miller & Bentlage (2024) describe stable insect-endosymbiont relationships that facilitate specialized metabolic functions. Transmission dynamics (vertical vs. horizontal) can vary with environmental conditions (Zchori-Fein *et al*., 2014), affecting seasonal symbiont retention. Thus, the observed variation is likely driven by a complex interplay of host biology, environmental stressors, and microbial ecology, warranting further investigation into seasonal symbiont-host interactions and their vectorial implications.

### Environmental Determinants of Endosymbiont Dynamics

Environmental variables showed strong correlations with endosymbiont prevalence. Sodalis showed the steepest decline under dry conditions, likely due to its facultative role and weaker host integration. In contrast, *Wolbachia* displayed intermediate sensitivity, potentially buffered by maternal transmission and tissue localization (Wernegreen, 2012). *Wigglesworthia*, as an obligate symbiont, maintained relatively high prevalence across conditions but still declined sharply above 32°C. These patterns reflect thermal sensitivities observed in other hematophagous insects (Ros *et al*., 2008; Roma *et al*., 2019), where elevated temperatures disrupt symbiont stability. The positive correlation with humidity supports findings by Smith *et al*. (2021) and Jones & White (2019), linking moist conditions with stable host physiology and symbiont survival. However, a non-linear decline at RH >80% suggests potential biological thresholds, possibly due to secondary stressors like fungal pathogens or altered host behavior (Haines *et al*., 2010). Despite adequate NDVI scores (∼0.65–0.72) during the wet season, prevalence still declined above 32°C, reinforcing the vulnerability of tsetse-symbiont systems to climatic extremes.

Comparatively, studies by Wernegreen (2012) and Guégan *et al*. (2020) have reported that endosymbionts such as *Buchnera* in aphids and *Wolbachia* in mosquitoes are highly susceptible to heat stress, leading to reduced densities or even symbiont loss under prolonged high temperatures. However, Bouyer *et al*. (2015) documented the persistence of tsetse flies at elevated temperatures (>30°C) in areas with adequate vegetation cover, which potentially buffered microclimatic extremes and moderated symbiont stress. Nonetheless, the non-linear decline at RH >80% in the current study suggests potential biological thresholds, possibly due to secondary stressors like fungal pathogens or altered host behavior (Haines *et al*., 2010). Despite adequate NDVI scores (∼0.65–0.72) during the wet season, prevalence still declined above 32°C, reinforcing the vulnerability of tsetse-symbiont systems to climatic extremes and suggesting narrower thermal margins for Nigerian *Glossina* populations compared to other regions.

### Implications for Targeted Vector Control and Climate Adaptation

The species- and site-specific ecological findings offer clear direction for tsetse control in northern Nigeria, *G. tachinoides’* widespread distribution suggests the effectiveness of wide-scale methods such as insecticide-treated cattle and mass trapping. The confinement of *G. palpalis palpalis* to humid zones necessitates focused control along water bodies. *Glossina morsitans submorsitans*, conservation of protected areas and flexible strategies addressing land-use change are crucial. On the microbial side, *Wolbachia’s* relative stability under moderate stress highlights its potential in paratransgenic control strategies, particularly in aridifying regions. However, the thermal and humidity sensitivities of all three symbionts point to possible challenges under future climate scenarios. Future research should prioritize predictive modeling that integrates environmental, host, and microbial variables to forecast changes in tsetse population dynamics and vector competence under projected climate conditions

### Climate Change Implications

The findings of this study underscore the susceptibility of tsetse fly populations and their endosymbionts to fluctuations in environmental conditions particularly temperature and humidity which are projected to become more erratic under climate change scenarios. Rising temperatures beyond critical thresholds (>32°C), as observed in this study, significantly reduced both *Glossina* abundance and endosymbiont prevalence. Such conditions could lead to geographic shifts in tsetse habitats, potential local extinctions in arid zones, and changes in disease transmission dynamics. Increased aridity and habitat fragmentation may compress the ranges of more climate-sensitive species like G. *morsitans submorsitans*, concentrating them in protected microclimates and reducing their overall contribution to trypanosome transmission. Conversely, the ecological plasticity of *G. tachinoides* may enable it to expand into newly suitable areas, potentially sustaining or even increasing transmission risks in unexpected locations.

Endosymbionts particularly *Sodalis* and *Wolbachia* demonstrated marked declines under thermal and desiccation stress, raising concerns about the long-term viability of symbiont-based control strategies such as paratransgenesis in warming environments. As endosymbionts are crucial for tsetse reproduction, immunity, and vector competence, their instability may impair host fitness or alter transmission potential.

Therefore, climate-resilient vector surveillance and control programs must incorporate microclimatic monitoring, ecological niche modeling, and thermal tolerance studies. This will be critical for anticipating changes in tsetse-endosymbiont dynamics and ensuring the sustainability of ongoing and future trypanosomiasis interventions.

## Conclusions

The ecological and microbiological dynamics of *Glossina* populations in northern Nigeria are shaped by complex interactions between species traits, environmental variables, and symbiotic relationships. The clear influence of climate on both tsetse abundance and endosymbiont stability underscores the need to integrate environmental surveillance into vector control planning. Future interventions must account for local habitat characteristics and climatic conditions to optimize outcomes. Broad-area control approaches may be effective for *G. tachinoides*, while targeted interventions are necessary for *G. palpalis palpalis* and *G. morsitans submorsitans*. Additionally, the use of endosymbiont-based strategies such as paratransgenesis should consider the ecological resilience of candidate symbionts under variable field conditions. As climate change continues to reshape vector ecologies, developing predictive models that incorporate temperature, humidity, vegetation, and host factors will be essential to sustaining control efforts and mitigating the risk of trypanosomiasis transmission in the region.

### Based on the findings of this study, the following recommendations are proposed

Integrated tsetse control in Nigeria should combine species-specific strategies with climate-sensitive planning. *G. tachinoides* responds to broad-area insecticide targets, while *G. palpalis palpalis* and *G. morsitans submorsitans* need localized approaches. Real-time surveillance, ecological modeling, and endosymbiont-based research are also essential

## Study Limitations

While this study offers valuable insights into the seasonal dynamics of *Glossina* species and their endosymbionts in northern Nigeria, certain limitations should be noted. First, the temporal scope was restricted to a single seasonal cycle, which may not account for inter-annual variability in tsetse abundance or symbiont prevalence. Second, the geographical coverage included only four ecological sites, which may limit the broader applicability of the findings to other tsetse-endemic areas in Nigeria. Finally, the reliance on stationary traps and their fixed placement could have introduced sampling bias, particularly affecting the capture of less mobile or more habitat-specific species such as *G. palpalis palpalis*.

## Open Data

10.6084/m9.figshare.29682023

## Authors’ Contributions

AA conceived the study. AA, EK and LSA designed the methodology. IHT, IUM, ZT, RAA, JA and UBM carried out field collections and laboratory analysis. AA and RGA conducted the statistical analysis and wrote the first draft. All authors reviewed and approved the final manuscript.”

## Ethics Approval

Not applicable. This study involved insect sampling which does not require ethical clearance.

